# Modeling the efficiency of filovirus entry into cells *in vitro*: Effects of SNP mutations in the receptor molecule

**DOI:** 10.1101/2019.12.19.882167

**Authors:** Kwang Su Kim, Tatsunari Kondoh, Yusuke Asai, Ayato Takada, Shingo Iwami

**Author notes:** Tatsunari Kondoh: Tsukuba Branch, WDB Co., Ltd., 1-6-1 Takezono, Tsukuba, Ibaraki, 305-0032, Japan. These authors contributed equally to this study (Kwang Su Kim, Tatsunari Kondoh). Correspondence and requests for materials should be addressed to: Shingo Iwami.

## Abstract

Interaction between filovirus glycoprotein (GP) and the Niemann-Pick C1 (NPC1) protein is essential for membrane fusion during virus entry. Some single-nucleotide polymorphism (SNPs) in two surface-exposed loops of NPC1 are known to reduce viral infectivity. However, the dependence of differences in entry efficiency on SNPs remains unclear. Using vesicular stomatitis virus pseudotyped with Ebola and Marburg virus GPs, we investigated the cell-to-cell spread of viruses in cultured cells expressing NPC1 or SNP derivatives. Eclipse and virus-producing phases were assessed by *in vitro* infection experiments, and we developed a mathematical model describing spatial-temporal virus spread. This mathematical model fit the plaque radius data well from day 2 to day 6. Based on the estimated parameters, we found that SNPs causing the P424A and D508N substitutions in NPC1 most effectively reduced the entry efficiency of Ebola and Marburg viruses, respectively. Our novel approach could be broadly applied to other virus plaque assays.

**Author Summary:** Ebola virus belongs to the family Filoviridae, together with Marburg virus and Cueva virus. In 2015, the World Health Organization included Ebola and Marburg viruses among the infectious diseases that should be globally prioritized because these filoviruses can cause severe hemorrhagic fever in humans and nonhuman primates, and antiviral agents to these viruses are very limited. Filovirus particles bear the envelope glycoprotein (GP), which is the only viral surface protein and thus responsible for receptor binding and membrane fusion. Interaction between filovirus GP and the Niemann-Pick C1 (NPC1) protein is essential for membrane fusion during virus entry. Some single-nucleotide polymorphism (SNPs) in two surface-exposed loops of NPC1 are known to reduce viral infectivity. However, the dependence of differences in entry efficiency on SNPs remains unclear. In this study, combining *in vitro* experiments and mathematical models, we evaluated on the interaction between GP and wildtype and mutant NPC1, enabling us to estimate the cellular entry efficiency during plaque formation.

## Introduction

In 2015, the World Health Organization (WHO) included Ebola (EBOV) and Marburg (MARV) viruses among the infectious diseases that should be globally prioritized. Some viruses of the family *Filoviridae*, which includes EBOV and MARV, cause severe hemorrhagic fever in humans and nonhuman primates. In recent years, more frequent filovirus outbreaks have been observed including multiple introductions of filoviruses into the human population, with important implications for worldwide public health [1]

Filovirus particles bear the envelope glycoprotein (GP), which is the only viral surface protein and thus responsible for receptor binding and membrane fusion [2]. Filovirus infection is initiated by binding of GP to attachment factors such as C-type lectins [3, 4], T-cell immunoglobulin and mucin domain 1 (TIM-1) and C-type lectins [5]. Virus particles are internalized into host cells via macropinocytosis and then delivered to late endosomes [6, 7]. GPs are proteolytically processed by cysteine proteases such as cathepsins B and L [8, 9]. This digested GP (dGP) can interact with the host endosomal fusion receptor, Niemann-Pick C1 (NPC1) protein, allowing fusion between the viral envelope and the host endosomal membrane [10, 11]. NPC1 is believed to be essential for filovirus entry into cells [12, 13].

Wang et al. showed that two surface-exposed loops of human NPC1 were important for interaction with dGP and that some amino acid substitutions in these loops reduced binding to dGP [14]. We previously investigated the potential effects of substitutions caused by naturally occurring single-nucleotide polymorphisms (SNPs) in these two loops and found that the P424A/D508N and S425L/D502E substitutions in human NPC1 reduced entry of vesicular stomatitis virus (VSV) pseudotyped with EBOV and MARV GPs, respectively [7]. We also found that the plaque sizes of replication-competent VSVs bearing EBOV and MARV GPs (VSVΔG-EBOV and VSVΔG-MARV) were reduced. However, it remains unclear how these SNPs and associated substitutions influence the spread of the viruses in plaque assays. In general, plaque formation is affected by multiple processes including cell-to-cell infection, viral production time, latent time, and intracellular replication. Because of this complexity, it is difficult to quantitatively understand the kinetics of viral infection.

In this study, we focused on the interaction between GP and NPC1, enabling us to estimate the cellular entry efficiency during plaque formation of VSVΔG-EBOV and VSVΔG-MARV. We employed viral infection assays combined with mathematical analyses as described previously [15–18] to quantitatively analyze viral entry efficiency and how this affected cell-to-cell spread. We found that the P424A and S425L substitutions reduced the entry efficiency of VSVΔG-EBOV by 47% and 21%, respectively, while the other SNPs and substitutions did not affect entry (reduction of <16%). Furthermore, we showed that our mathematical model recapitulates the process of merging viral plaques. This method could also be applied to plaque assays for other viruses and could be used to improve *in vitro* determination of the effects of mutations on viruses and target cells.

## Results

### Distribution of infectious phases of VSVΔG-EBOV and VSVΔG-MARV

We assumed that amino acid mutations of the cellular NPC1 protein only changed virus entry efficiency; the eclipse phase and infectious phase remained unchanged. To estimate the infectious phase for VSVΔG-EBOV and VSVΔG-MARV, we performed virus production assays using Vero E6/NPC1-KO cells expressing human NPC1 (293T-NPC1) (**Fig.1A** and **B**, left panels; see also **Methods**). In three of the four experiments (Exp1, 3, and 4), VSVΔG-EBOV-inoculated cells started to produce infectious virus particles at 6 h post inoculation. In Exp2, the inoculated cells started to produce virus at 9 h post-inoculation. The virus-producing cells died by 33 h post-inoculation in all experiments. Thus, we assumed that infected cells which produce infectious viruses are in “infectious phase”. In our experiments, since the discrepancy in virus production time was 3 h and the exact virus production time was unknown, candidate groups were classified as follows: candidates for the time when infected cells begin to produce virus were 4, 5, and 6 h (occurring in three instances), while candidates at 7, 8, and 9 h occurred only once. In all four experiments, the death of all infected cells was observed at 33 h post-inoculation. This suggested that the candidates for time of cell death were 31, 32, and 33 h, and if we consider the above virus production start time, the infectious phase was between 22 and 29 h in duration. By contrast, for VSVΔG-MARV-inoculated cells, the infectious phase was between 25 and 29 h in duration.

**Figure 1.**
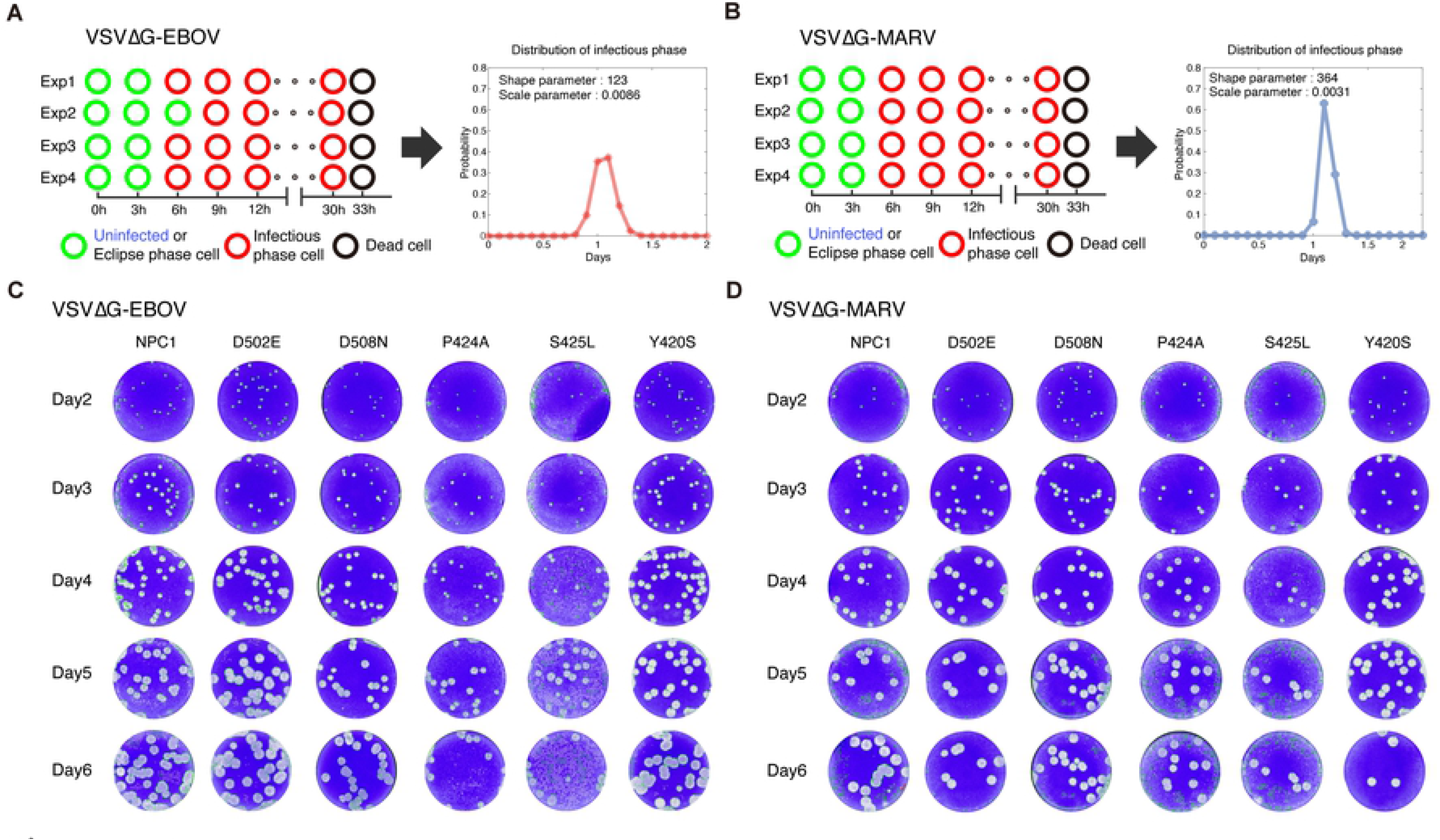
Virus production assays and plaque assays: Virus production assays using Vero E6/NPC1-KO cells expressing human NPC1. Four independent experiments recording cell state every 3 h for VSVΔG-EBOV and VSVΔG-MARV are shown in the left panels of **(A)** and **(B)**, respectively. Estimated distributions of the infectious phases of each are is also shown in the right panel. Plaque formation of VSVΔG-EBOV **(C)** and VSVΔG-MARV **(D)** on Vero E6 cells expressing wildtype NPC1 and five SNP mutants on days 2-6 are shown. Gray spots represent plaques formed by dead cells.

Because the infectious phase was relatively long compared with the duration of the eclipse phase, we assumed that the infectious phase follows the Erlang distribution as described previously [19, 20]. The equivalence between the expression for, and the parameters of, the probability density functions of the Erlang distributions is shown as follows [21]:

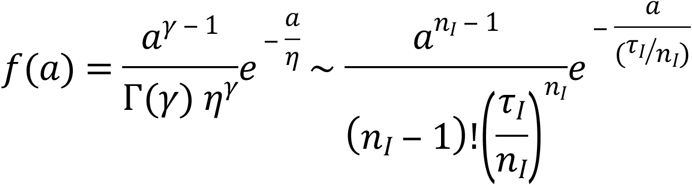

The shape (*γ* = *n*_*I*_ = 123 and 364) and scale 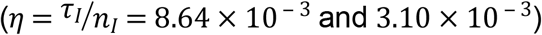 parameters of the Erlang distribution for VSVΔG-EBOV and VSVΔG-MARV were estimated by fitting the candidates for the infectious phase, respectively (**Table 1**). Note that *n*_*I*_ and *τ*_*I*_ correspond to the number of “subdivided” compartments and the average duration of the infectious phase, respectively, in Eq.(4) [20, 21] (see later). The probability density functions of the estimated Erlang distributions of infectious phases are shown in the right panels of **Fig.1A** and **B**. Conversely, we assumed that the eclipse phase with relatively short duration follows an exponential distribution as in many basic virus dynamics models [22–24]. We planned to quantify the eclipse phase for VSVΔG-EBOV and VSVΔG-MARV from the results of plaque assays together with other parameters (see later).

**Table 1.** Parameter values estimated from plaque assay and virus production assay.

| Parameter Name | Symbol | Unit | Virus type | Wildtype | D502E | D508N | P424A | S425L | Y420S |
| --- | --- | --- | --- | --- | --- | --- | --- | --- | --- |
| Number of subdivided compartments | $n_I$ | --- | VSVΔG-EBOV | | | 123 | | | |
| | | | VSVΔG-MARV | | | 364 | | | |
| Length of infectious phase | $\tau_I$ | day | VSVΔG-EBOV | | | 1.06 | | | |
| | | | VSVΔG-MARV | | | 1.13 | | | |
| Rate constant for infections | $\omega$ | (cell · day)<sup>-1</sup> | VSVΔG-EBOV | | | 0.38 | | | |
| | | | VSVΔG-MARV | | | 0.41 | | | |
| Length of eclipse phase | $1/k$ | day | VSVΔG-EBOV | | | 0.19 | | | |
| | | | VSVΔG-MARV | | | 0.22 | | | |
| Initial radius of viral plaque | $r_0$ | mm | VSVΔG-EBOV | | | 0.23 | | | |
| | | | VSVΔG-MARV | | | | | | |
| Relative virus entry efficiency | $\alpha$ | --- | VSVΔG-EBOV | 1.00 | 1.26 | 0.84 | 0.53 | 0.79 | 1.05 |
| | | | VSVΔG-MARV | 1.00 | 1.12 | 0.93 | 1.1 | 1.05 | 1.15 |

### Spatial-temporal mathematical model for viral plaque amplification

To quantify and compare the filovirus entry efficiency among cells expressing wildtype and SNP-mutant NPC1, we performed viral plaque assays using cells expressing wildtype NPC1 (293T-NPC1) and five SNP mutants for VSVΔG-EBOV and VSVΔG-MARV (**Fig.1C** and **D**). The viral plaque is considered as the area formed by dead cells, and thus the plaque radius is defined by the distance from the center of plaque to its edge. We measured the average sizes of independent plaques (see **Methods**) and used them to quantify spatial-temporal VSVΔG-EBOV and VSVΔG-MARV spread. First, we developed a novel mathematical model for viral plaque amplification as follows. Because monolayers of cells were overlaid with agar media in our plaque assay, there was no cell movement. Only cell-to-cell infections between infected and adjacent uninfected cells occur. We assumed that the inner infected cell infects only adjacent outer target cells (**Fig.2A**). To describe the infection dynamics of virus in the plate, we derived the following mathematical model including two independent variables (time *t* and radius of circle *r*) and four state variables (uninfected, eclipse phase, infectious phase and dead cells):

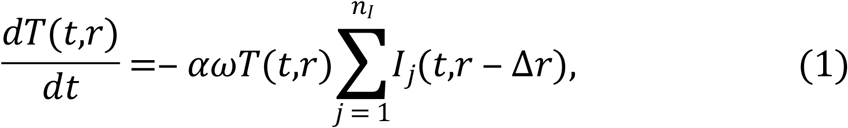

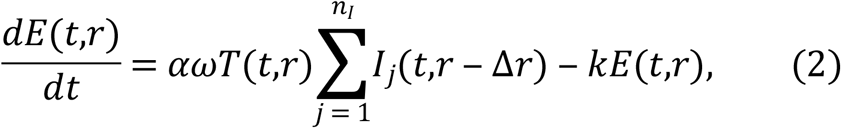

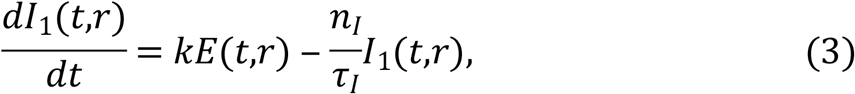

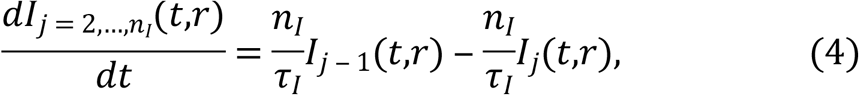

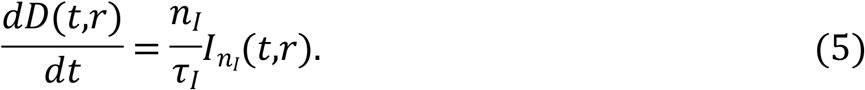

**Figure 2.**
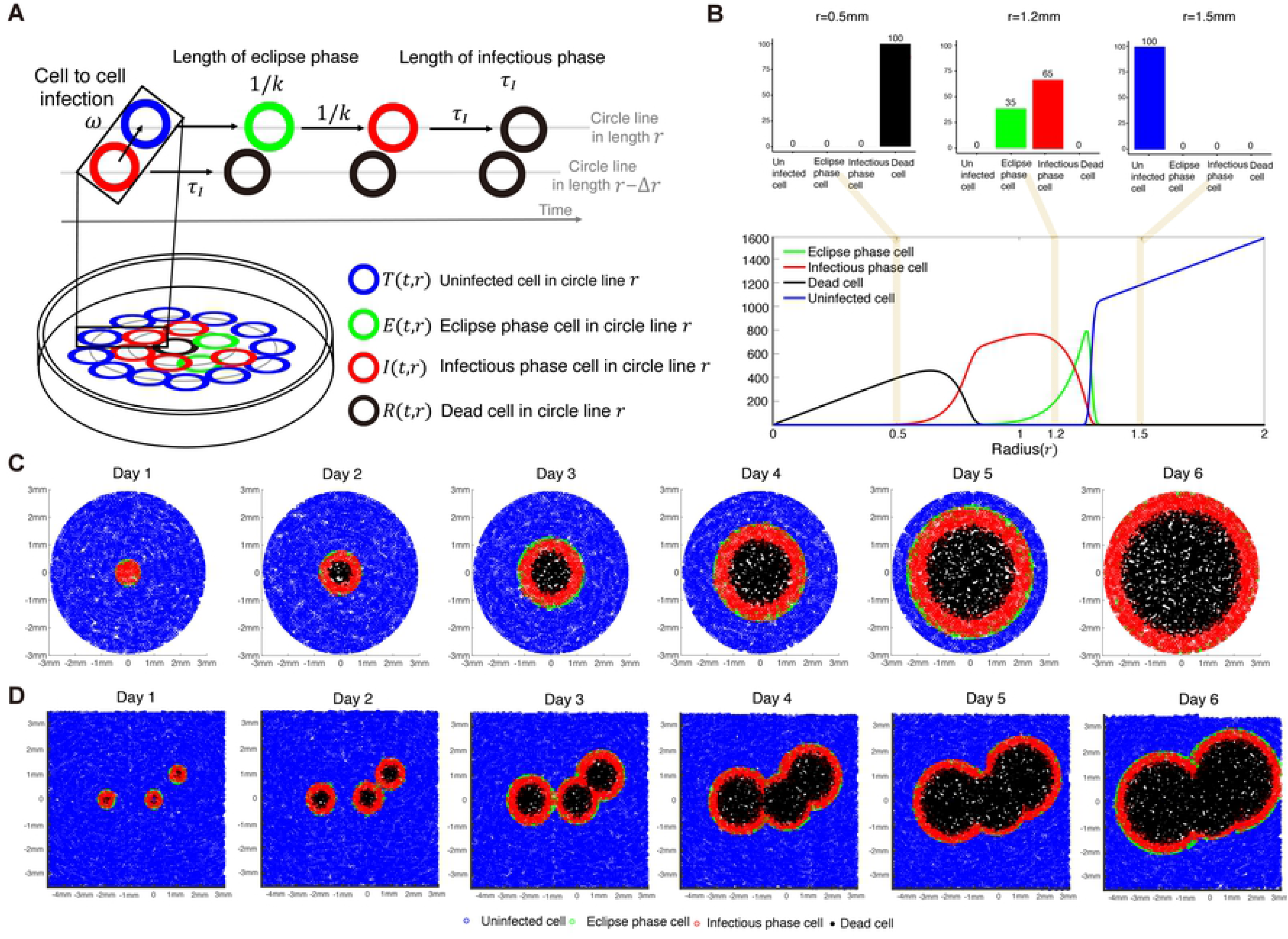
Modeling and visualizing viral plaque amplification: Modeling spatial-temporal dynamics of viral plaque amplification is shown in **(A)**. A representative simulation for plaque amplification on day 3 is shown in **(B)**. The ratio of cells for each radius (e.g., 0.5mm, 1.2mm and 1.5mm) can be calculated in the top panels. In **(C)**, a representation of a single viral plaque amplification is described for each day. Merging of three plaques can be also described by considering overlapping portions over time in **(D)**.

The initial conditions were: *I*_1_(0,*r*) = 2*π* × *r*/0.008 for *r* ≤ *r*_0_ and 0 for *r* > *r*_0_, *T*(0,*r*) = 0 for *r* ≤ *r*_0_ and 2*π* × *r*/0.008 for *r* > *r*_0_, and *E*(0,*r*) = *D*(0,*r*) = 0 for *r* > 0. Here *T*(*t*,*r*), *E* (*t*,*r*), *I*_*j*_(*t*,*r*) and *D*(*t*,*r*) represent the numbers of uninfected, eclipse phase, infectious phase and dead cells, respectively. Note that Eq. (4) is derived from the following integro-differential equation by “linear-chain-trick” which is discussed in detail elsewhere [25–27]:

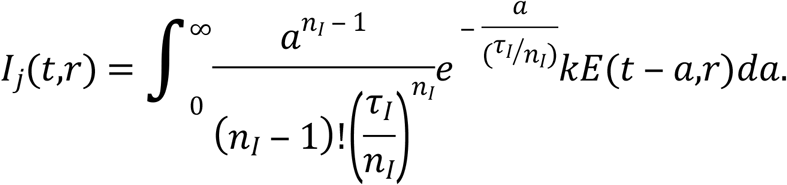

We also assumed that infectious cells of the *I*_1_ compartment were inoculated only in a radius of less than *r*_0_ and there were no other infected cells at the initial time. The parameters *ω* and 1/*k* represent the infection rate of cells expressing 293T-NPC1 and the length of the eclipse phase, respectively, and thus *α* is the relative virus entry efficiency into target cells bearing amino acid mutations in the cellular NPC1 (i.e., we fixed *α* = 1 for 293T-NPC1-expressing cells). Values of entry efficiency (*α*) larger than 1 means shorter the virus entry time and more efficient infection of uninfected cells.

### Simulating and visualizing viral plaque amplification

The number of target cells in our plate was initially distributed as follows. The radius of the plate was 17.35 mm and the average cell radius was 0.004 mm. This implies that there are 17.35 mm/0.008 mm=2169 circle lines in which cells are distributed in the plate. If the circumference of the circle line is divided by the diameter of the cell, we can obtain the number of cells distributed in one circle line of radius *r*, that is, 2*πr*/0.008. Since the radius of each circle line increases proportionally to the cell interval (0.008 mm), total cell number in the plate can be calculated as 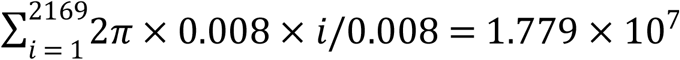, and was consistent with the number of cells used in our experiments.

Using a finite difference method, we computed a numerical solution of Eqs. (1-5) with respect to time *t* and circle radius *r* in the plate. In the bottom panel of **Fig.2B**, for example, the distribution of each cell according to the radius at day 3 is shown. Dead cells (black line) are located at a radius of 0.8 mm from the center of the plaque which is considered as “a plaque radius” in our simulation. That is, we defined the simulated viral plaque radius as the length of a plaque from its center to the edge of dead cells in the plaque. Here the edge is defined as the first circle line which does not include any dead cell (dead cells fewer than “1” were not counted). Infectious cells (red line) are distributed from 0.5 to 1.3 mm in radius, and eclipse phase cells (green line) gradually increase as the radius increases, peaking at 1.3 mm. This differs from the distribution of infectious cells in the wide radius region because the time spent in the infectious phase is relatively longer than the time spent in the eclipse phase. Although we assumed any cells in an inner circle line can interact with those in the adjacent outer circle line, our mathematical model, Eqs. (1-5), can describe virus amplification in the plaque which is a minimum model for quantitatively analyzing “plaque size” (see later).

Next, to visualize amplification of an average viral plaque, we calculated the percentage of cells corresponding to each state with respect to time *t* and circle radius *r* based on Eqs. (1-5). For example, in the top panels of **Fig.2B**, the corresponding percentage of each cell at radius 0.5 mm, 1.2 mm and 1.5 mm on day 3 is shown. There were only dead cells at radius 0.5 mm. At radius 1.2 mm (depicted in the second bar graph), eclipse phase and infectious phase cells comprised 35% and 65% of cells, respectively. Only uninfected cells were present at radius 1.5 mm, as infection had not yet occurred. After calculating the ratio of cells at each radius, we chose each cell state depending on the ratio at a circle line by line. By simulation over time, in **Fig.2C**, we show a representation of viral plaque amplification. On day 1, no plaque has yet been generated; only uninfected cells, infectious cells and eclipse phase cells are distributed. On day 2, a plaque of dead cells is forming (black circle). The eclipse phase cells, which are represented by a green circle, are narrowly distributed on the edge of the infectious cell area, and the infectious cells are in turn distributed in the wide radius region because the period spent in the infectious phase is longer than that in the eclipse phase as explained above. Note that in **Fig. 2C**, we show only the change of composition within the area inside a 3-mm radius, not the whole plate. All uninfected cells disappear inside this radius, and only infectious cells and dead cells remain at day 6.

In the plaque assay data for day 6 in **Fig.1CD**, we can see that several plaques expanded to form a larger contiguous plaque. We demonstrated that Eqs. (1-5) can reproduce these merging plaques (see detail in **Supplementary Note**). In **Fig. 2D**, the dynamics of the merging three plaques located at different positions are shown as snapshots. In **Supplementary Movie 1** and **2**, we also showed how two and three plaques merge in a spatial-temporal manner.

### Quantifying filovirus entry efficiency for cellular NPC1 SNP mutations

The plaque radii generated by infection of wildtype and mutant NPC1-expressing cells were measured and compared with fitting results of the radius of viral plaques simulated by Eqs. (1-5) for the six amino acid mutations in both viruses, as shown in **Fig.3A** (see also **Methods**). For all of the NPC1 mutants, the plaque radius for VSVΔG-EBOV was smaller than that for VSVΔG-MARV, meaning that the infectivity (i.e., entry efficiency) of VSVΔG-MARV was greater than that of VSVΔG-EBOV, regardless of the sequence of NPC1. The estimated parameter values are shown in **Table 1** and the viral entry efficiency, *α*, is shown in **Fig.3B**. In the case of VSVΔG-MARV, the entry efficiencies into wildtype and mutant NPC1-expressing cells were similar. Indeed, the entry efficiencies of NPC1 mutants were slightly greater than that of 293T-NPC1 except for the D508N and Y420S substitutions, which had the highest entry efficiency (15% above wildtype). In the case of VSVΔG-EBOV, the P424A, S425L, and D508 substitutions in NPC1 resulted in lower entry efficiencies than 293T-NPC1. The P424A substitution showed the lowest entry efficiency (47% reduction compared with 293T-NPC1). In contrast, the D502E substitution, which had the highest entry efficiency among the NPC1 mutants for VSVΔG-EBOV, demonstrated 26% higher entry efficiency than wildtype NPC1 (discussed below).

**Figure 3.**
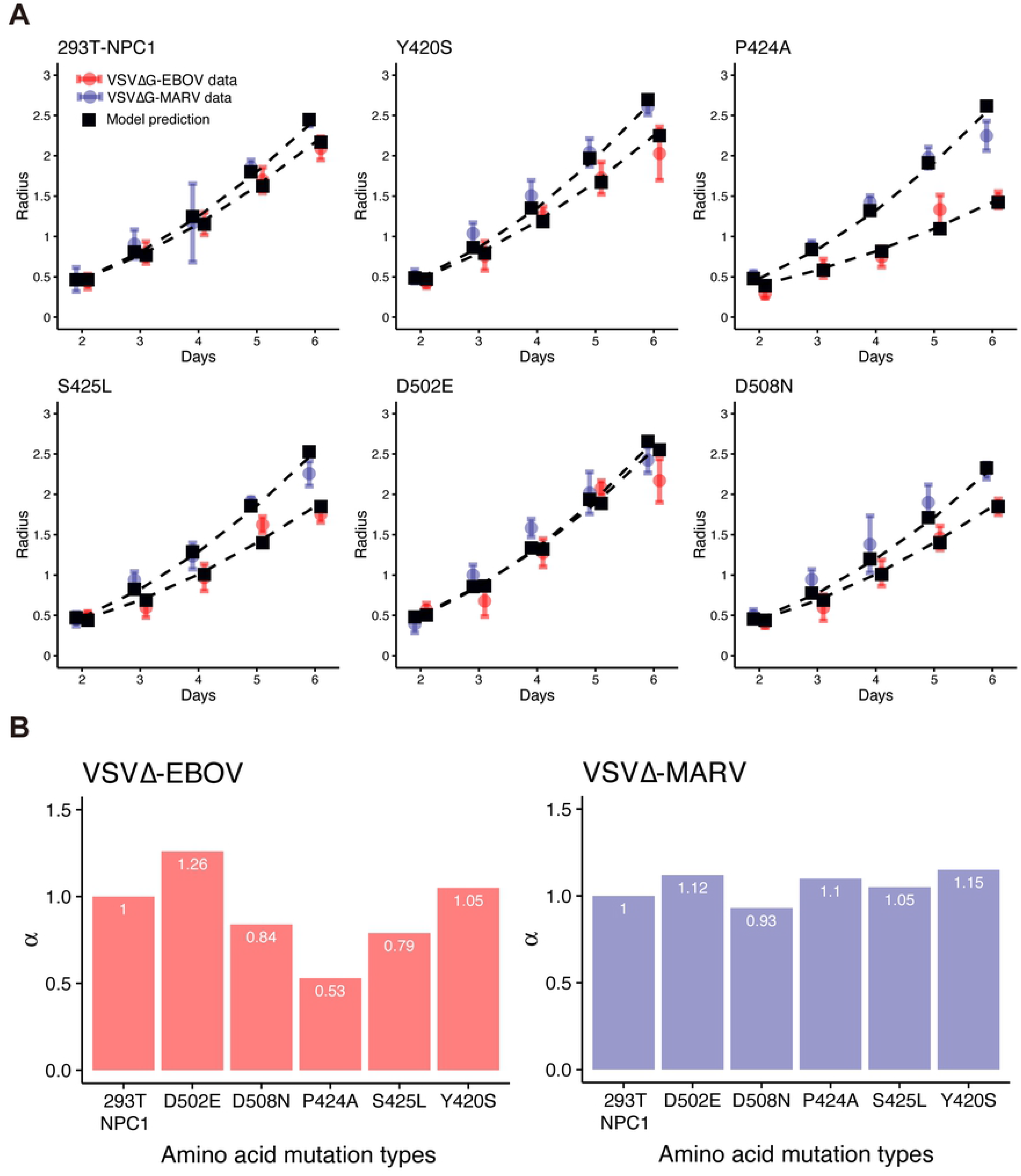
Quantifying VSVΔG-EBOV and VSVΔG-MARV entry efficiency for different NPC1 SNPs: Fits of the mathematical model, Eqs. (1–5), to the experimental data of VSVΔG-EBOV and VSVΔG-MARV in plaque assays are shown in **(A)**. Black squares represent plaque radii from simulations with best-fit parameters for each SNP. Blue and red circles and bars represent the means and standard deviations of the plaque radii following VSVΔG-EBOV and VSVΔG-MARV infection, respectively, on wildtype and mutant NPC1-expressing cells. Compared with infection of 293T-NPC1-expressing cells by VSVΔG-EBOV and VSVΔG-MARV, relative entry efficiency, *α*, fitting the plaque radius dataset was estimated for each NPC1 SNP and shown in **(B)**.

## Discussion

In general, plaque formation is affected by multiple processes including viral entry, membrane fusion, genome replication, transportation, and virion assembly efficiency. Because of this complexity, it is difficult to quantitatively understand the kinetics of viral infection and how the efficiency of entry is affected by individual SNPs. Moreover, the time course of plaque data is challenging to understand intuitively. Additionally, there is no experimental technique available to measure only entry efficiency as an absolute value excluding other causes of viral spread. To address this point, we employed two viral infection assays (i.e., plaque-forming assay and virus production assay) combined with mathematical analyses to quantitatively analyze how particular SNPs affected virus entry. Previous mathematical models which describe plaque expansion considered the diffusion of virus [28–31], but under our experimental conditions, the infected cells were overlaid with agar and thus only cell-to-cell infection was monitored. Agent-based models can describe spatially explicit mechanisms [32, 33]. However, these approaches are difficult to directly fit to plaque radius data in a time course manner. A model including the infection term for the decreasing proportion of cells contributing to cell-to-cell was also suggested [34].

We developed a simple but well approximated mathematical model, i.e., Eqs. (1-5), that can analyze plaque assay data with minimal assumptions to quantitatively compare and analyze the virus entry efficiency by focusing on the interactions between GP and NPC1. VSVΔG-EBOV and VSVΔG-MARV usually grow in cultured cells as well as VSV (taking several hours to produce cytopathic effect (CPE)), whereas EBOV and MARV do not grow as rapidly as VSV. These pseudotyped VSVs enabled us to concentrate on the interaction between GP and NPC1 without considering other factors, since all other viral proteins are identical between VSVΔG-EBOV and VSVΔG-MARV. Thus, we could evaluate entry efficiency and objectively compare each SNP after parameterization by Eqs. (1-5) for VSVΔG-EBOV and VSVΔG-MARV. The entry efficiencies of VSVΔG-MARV in cells expressing NPC1 with different SNPs and substitutions ranged from −7% to +15% compared with 293T-NPC1 (wildtype human NPC1), while the efficiencies of VSVΔG-EBOV showed relatively large changes ranging from −47% to +26%. These results indicated that the entry efficiency of VSVΔG-EBOV was more sensitive to changes in NPC1 sequence. Notably, the P424A and S425L substitutions reduced the entry efficiency of VSVΔG-EBOV by 47% and 21%, respectively, but only reduced the entry efficiency of VSVΔG-MARV by 10% and 5%, respectively. This supports the possibility that the P424A and S425L substitutions have different effects due to differences between Ebola and Marburg viruses observed in previous studies [35]. The D508N substitutions reduced entry efficiency for both Ebola and Marburg viruses. We found that P424A and D508N substitutions significantly reduced the entry of VSVΔG-EBOV. Although it might be difficult to completely apply the values determined here for entry efficiency to *bona fide* EBOV and MARV infection, our result is consistent with previous investigations [35]. We highlight that pseudotyped viruses are useful in mathematical model-based quantitative data analyses focusing on a specific molecular interaction. For example, if we polymerase mutations into EBOV and MARV, similar approach might reveal the differences in viral replication efficiency and their dependence on mutations. Our novel approach could be broadly applied to other virus plaque assays.

Through this experimental-mathematical investigation, we quantified the entry efficiency of VSVΔG-EBOV and VSVΔG-MARV based on cell-to-cell spread during plaque formation and found differences among cells bearing SNPs and amino acid substitutions in the filovirus receptor (i.e., NPC1). Although there have been some studies of asymptomatic filovirus infection [36, 37], the mechanisms through which individuals appear to be inherently resistant to EBOV and MARV have not yet been understood. It will be of interest to investigate NPC1 variation and its influence on EBOV and MARV entry efficiency and also to identify genetic backgrounds that affect the susceptibility of humans to filovirus infection, both of which will provide important information for understanding filovirus disease progression and host restriction. Combining *in vitro* experiments and mathematical models gradually provides detailed quantitative insights into the kinetics of virus infection [17, 38, 39]. Thus, our method may also be applied to understanding the roles of genetic polymorphisms in human susceptibility to filoviruses.

## Methods

### Viruses and cells

Replication-competent recombinant VSVs pseudotyped with EBOV (Mayinga) and MARV (Angola) GPs (VSVΔG-EBOV and VSVΔG-MARV, respectively) were generated as described previously [40]. VSVΔG-EBOV and VSVΔG-MARV were propagated in Vero E6 cells and stored at −80°C until use. Infectivity of the viruses in each cell line was determined by a plaque-forming assay as described previously [41]. All work using these viruses was performed in the BSL-3 laboratories at the Research Center for Zoonosis Control, Hokkaido University, Japan. Vero E6 cells (ATCC^®^ CRL-1586™), NPC1-knockout Vero E6 (Vero E6/NPC1-KO), and Vero E6 cell lines stably expressing each NPC1 SNP (293T-NPC1, Y420S, P424A, S425L, D502E, D508N) [7] substitution were grown in Dulbecco’s modified Eagle’s medium (DMEM, Sigma) supplemented with 10% fetal calf serum (FCS).

### Plaque assay

VSVΔG-EBOV and VSVΔG-MARV (multiplicity of infection, MOI = 0.0005 in Vero E6 cells) were inoculated onto monolayers of each cell line in six-well tissue culture plates (Corning). After adsorption for 1 h, the inoculum was completely removed, and the cells were overlaid with Eagle’s minimal essential medium containing 1.0% Bacto Agar (BD) and then incubated for 2–6 days at 37°C. Cells were stained with 0.5% crystal violet in 10% formalin at 24 h intervals. Plaque images in the wells were captured and each plaque size (mm^2^) was measured using a CTL-ImmunoSpot^®^ S6 Macro Analyzer equipped with ImmunoCapture ver. 6.5 and BioSpot 5.0 software (Cellular Technology Ltd. USA). We examined all plaques which were completely separated from one another. Average sizes of independent plaques were used for mathematical model-based quantitative data analyses.

### Virus production assay

Vero E6 cells grown in 96-well tissue culture plates (Corning) were inoculated with VSVΔG-EBOV and VSVΔG-MARV (MOI = 1.0). After adsorption for 1 h, the inoculum was completely removed. One hundred microliters of growth medium (DMEM supplemented with 10% FCS) were added into each well and then incubated for 33 h at 37°C. Supernatants of the culture medium were collected at 3 h intervals and frozen at −80°C until use. To check the presence of infectious virus particles in the collected supernatants, confluent monolayers of Vero E6 cells on 96-well tissue culture plates (Corning) were inoculated with the supernatant collected at each time point. After incubation for 3 days at 37°C, virus infection was assessed by the presence of CPE. This assay enabled us to predict the time when infected cells shift from non-virus-producing to virus-producing cells (i.e., eclipse phase).

### Data fitting and parameter estimation

Because the number of experimental measured plaques and the radius of each plaque were different (we simply employed the mean radius of plaques which had not merged), we used in our data fitting the weighted least square method considering the means and standard deviations of the observations, 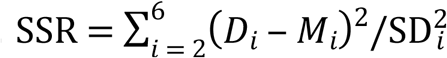, where *D*_*i*_ and SD_*i*_ are the mean and standard deviation of the plaque radius in experiments, respectively, and *M*_*i*_ is the radius of the plaque in our simulation at day *i* = 2,3,…,6. Using estimated *τ*_*I*_ and *n*_*I*_ in the virus production assay, we estimated the parameters *ω* and *k* for VSVΔG-EBOV and VSVΔG-MARV, and the common initial value of *r*_0_ from the plaque assay with 293T-NPC1. With these estimated parameters, we quantified the viral entry efficiency, *α*, for VSVΔG-EBOV and VSVΔG-MARV from the plaque assay with amino acid mutations in NPC1 (D502E, D508N, P424A, S425L and Y420S). All estimated parameters are summarized in **Table 1** and **Fig.4B**.

## Acknowledgments

This study was supported in part by the Basic Science Research Program through the National Research Foundation of Korea funded by the Ministry of Education (2019R1A6A3A12031316 to K.S.K.); Grants-in-Aid for Scientific (KAKENHI) Scientific Research B (18KT0018, 18H01139, and 16H04845 to S.I.), by Scientific Research on Innovative Areas (19H04839 and 18H05103 to S.I.); AMED CREST (19gm1310002 to S.I.); AMED J-PRIDE (19fm0208006s0103, 19fm0208014h0003, and19fm0208019h0103 to S.I.); the AMED Research Program on HIV/AIDS 19fk0410023s0101 (to S.I.); the Research Program on Emerging and Re-emerging Infectious Diseases (19fk0108050h0003 to S.I.); the Program for Basic and Clinical Research on Hepatitis (19fk0210036h0502 to S.I.); the Program on the Innovative Development and the Application of New Drugs for Hepatitis B (19fk0310114h0103 to S.I.); JST MIRAI (to S.I.); JST CREST (to S.I.); Mitsui Life Social Welfare Foundation (to S.I.); Shin-Nihon of Advanced Medical Research (to S.I.); Suzuken Memorial Foundation (to S.I.); Life Science Foundation of Japan (to S.I.); SECOM Science and Technology Foundation (to S.I.); The Japan Prize Foundation (to S.I.); Toyota Physical and Chemical Research Institute (to S.I.); and Fukuoka Financial Group, Inc. (to S.I.). We thank Edanz Group (https://en-author-services.edanzgroup.com/) for editing a draft of this manuscript.

## Authors’ contributions

Conceived and designed the study: SI. Analyzed the data: KSK, SI. Carried out the experiments: TK, AT. Wrote the paper: KSK, SI, TK, AT. All authors read and approved the final manuscript.

